## Supplementary Data 3 for "Notebook-based alignment of human and agentic reasoning in single-cell biology"

### Supplementary Data 3: Prompts

In all cases, "Output a Jupyter notebook in json format to address the following:" was added to the prompt for the reference model but not for *kai* because *kai's* internal prompts already provide for this guidance.

**Table of Contents**


***Scenario 1:*** ... **1**

**PBMC:** ... 1

**Breast cancer:** ... 1

**Lung:** ... 2

***Scenario 2:*** ... **2**

**Neutral prompt:** ... 2

**Weakly biased prompt:** ... 3

**Strongly biased prompt:** ... 3

***Scenario 3:*** ... **4**

**Question prompt** ... 4

**Execution prompt** ... 4


**Scenario 1:**

*PBMC:*

Output a Jupyter notebook in json format to address the following:

Use tutorials on the following concepts:

- Basic celltypist usage from the celltypist repository.
- Batch effects & harmony from scverse if available.

Perform a leiden clustering to use as a basis for cell type annotation. Show the downloaded models in celltypist (don't download any models). Based on that list and any information that you can find about these models, choose two that are applicable to this tissue. Use those models to annotate the leiden clusters with cell type labels. Then, assemble a marker gene panel from literature sources for the cell types predicted by the models, matching their cell type granularity, taking care to assemble a comprehensive panel with several genes per cell type. Use that panel to compute gene expression scores for each cell type. Interpret these scores per cluster and use these scores to assign cell type labels to clusters. Conclude to what degree one can use the results from this celltypist analysis for downstream analyses by working through the agreement of celltypist with the marker gene approach - take care to account for differences in cell type naming and granularity between the different annotations. Generate one final cell type annotation for downstream analyses.

<u>*Lung:*</u>
Output a Jupyter notebook in json format to address the following:

You are given single-cell RNA-seq data from human lung tissue of a COVID-19 patient as adata. Cells are already filtered and ready for analysis. Gene expression data is given as raw counts in adata.layer["counts"].X and as log1p-normalized in adata.X.

Use tutorials on the following concepts:
- Basic celltypist usage from the celltypist repository.
- Basic single-cell analysis tutorial for leiden clustering.

Perform a leiden clustering to use as a basis for cell type annotation. Show the downloaded models in celltypist (don't download any models). Based on that list and any information that you can find about these models, choose two that are applicable to this tissue. Use those models to annotate the leiden clusters with cell type labels. Then, assemble a marker gene panel from literature sources for the cell types predicted by the models, matching their cell type granularity, taking care to assemble a comprehensive panel with several genes per cell type. Use that panel to compute gene expression scores for each cell type. Interpret these scores per cluster and use these scores to assign cell type labels to clusters. Conclude to what degree one can use the results from this celltypist analysis for downstream analyses by working through the agreement of celltypist with the marker gene approach - take care to account for differences in cell type naming and granularity between the different annotations. Generate one final cell type annotation for downstream analyses.

**Scenario 3:**
Each run started with a chat query to kai or the reference models and was followed with an execution query once the models had defined their research question in response to the first question.
