## Supplementary Figures for "Notebook-based alignment of human and agentic reasoning in single-cell biology"

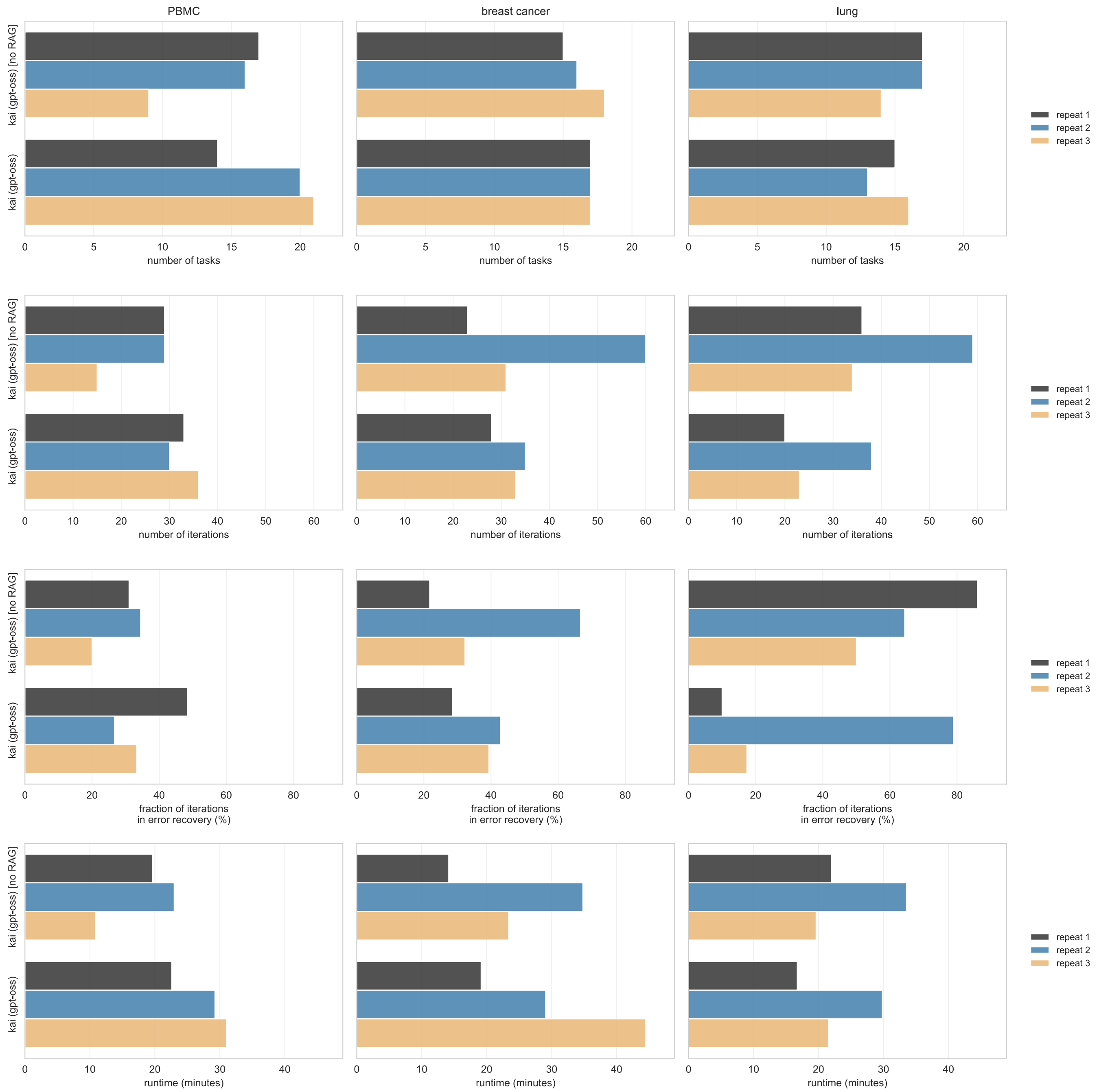


**Supp. Fig. 1: kai run characteristics in the cell type annotation scenario.** Shown are from top to bottom the number of tasks in the task list, the number of iterations performed by *kai*, the fraction of iterations in error recovery and the total run time of *kai* in minutes for each tissue in the cell type annotation scenario (PBMC, breast cancer and lung from left to right), shown for each repeat.

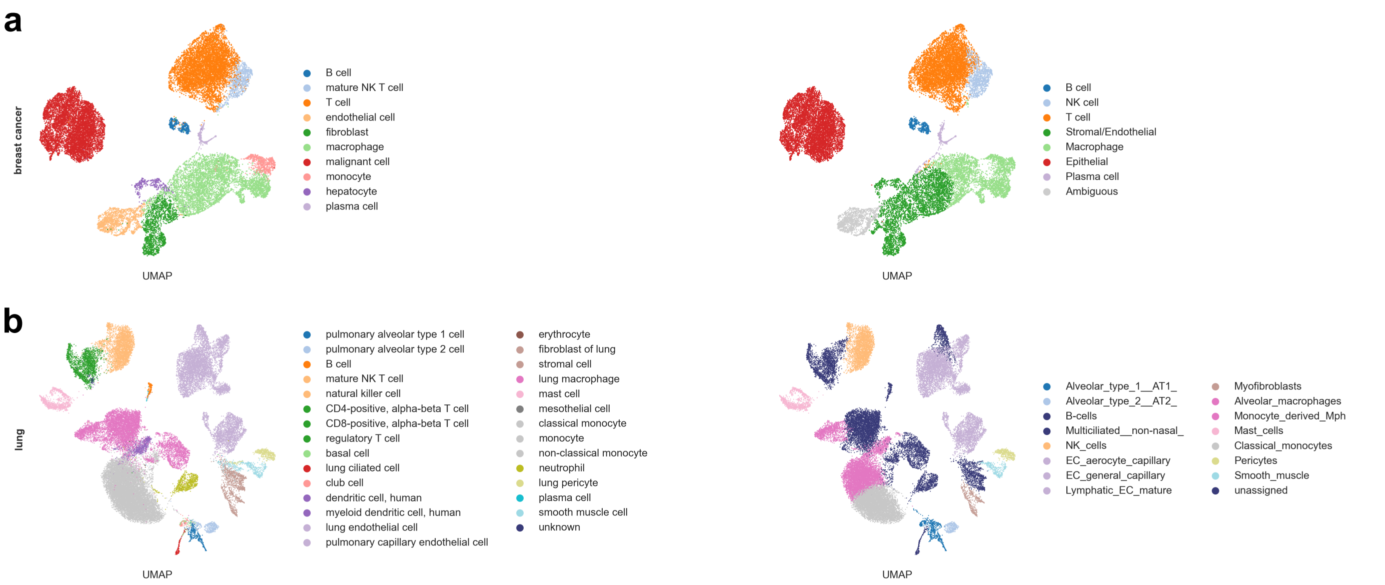


**Supp. Fig. 2: Further cell type annotation cases.** Uniform manifold approximation and projection (UMAP) of the breast cancer (**a**) and lung (**b**) dataset with the ground truth (left) and predicted labels (right) of one kai model run. Note that colors are chosen to match across panels so that multiple labels received the same color in some cases, as documented in the legends. This figure extends **Fig. 1d,e**.

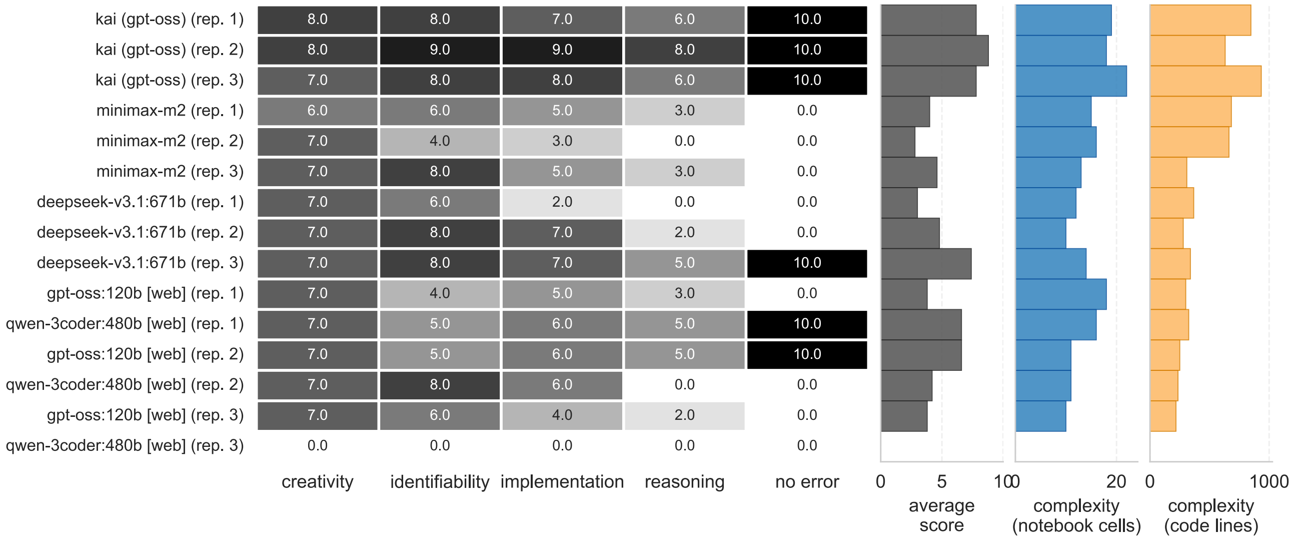


| | creativity | identifiability | implementation | reasoning | no error |
|---|---|---|---|---|---|
| kai (gpt-oss) (rep. 1) | 8.0 | 8.0 | 7.0 | 6.0 | 10.0 |
| kai (gpt-oss) (rep. 2) | 8.0 | 9.0 | 9.0 | 8.0 | 10.0 |
| kai (gpt-oss) (rep. 3) | 7.0 | 8.0 | 8.0 | 6.0 | 10.0 |
| minimax-m2 (rep. 1) | 6.0 | 6.0 | 5.0 | 3.0 | 0.0 |
| minimax-m2 (rep. 2) | 7.0 | 4.0 | 3.0 | 0.0 | 0.0 |
| minimax-m2 (rep. 3) | 7.0 | 8.0 | 5.0 | 3.0 | 0.0 |
| deepseek-v3.1:671b (rep. 1) | 7.0 | 6.0 | 2.0 | 0.0 | 0.0 |
| deepseek-v3.1:671b (rep. 2) | 7.0 | 8.0 | 7.0 | 2.0 | 0.0 |
| deepseek-v3.1:671b (rep. 3) | 7.0 | 8.0 | 7.0 | 5.0 | 10.0 |
| gpt-oss:120b [web] (rep. 1) | 7.0 | 4.0 | 5.0 | 3.0 | 0.0 |
| qwen-3coder:480b [web] (rep. 1) | 7.0 | 5.0 | 6.0 | 5.0 | 10.0 |
| gpt-oss:120b [web] (rep. 2) | 7.0 | 5.0 | 6.0 | 5.0 | 10.0 |
| qwen-3coder:480b [web] (rep. 2) | 7.0 | 8.0 | 6.0 | 0.0 | 0.0 |
| gpt-oss:120b [web] (rep. 3) | 7.0 | 6.0 | 4.0 | 2.0 | 0.0 |
| qwen-3coder:480b [web] (rep. 3) | 0.0 | 0.0 | 0.0 | 0.0 | 0.0 |

**Supp. Fig. 3: Detailed model scores per run in the open question scenario.** This figure extends **Fig. 3a** by showing the exact scores per model run rather than average with error bars. Evaluation of *Jupyter notebooks* addressing the open-ended question scenario by kai and reference LLMs operated in one-shot mode. Scores were assigned by a reviewer agent to each notebook to evaluate creativity of the question, feasibility of addressing it on the given dataset, implementation, and reasoning over the resulting findings (**Methods**). number of errors: number of notebooks out of three repeats that terminated with error; notebook cells/lines of code: complexity measures on the generated *Jupyter notebooks*.

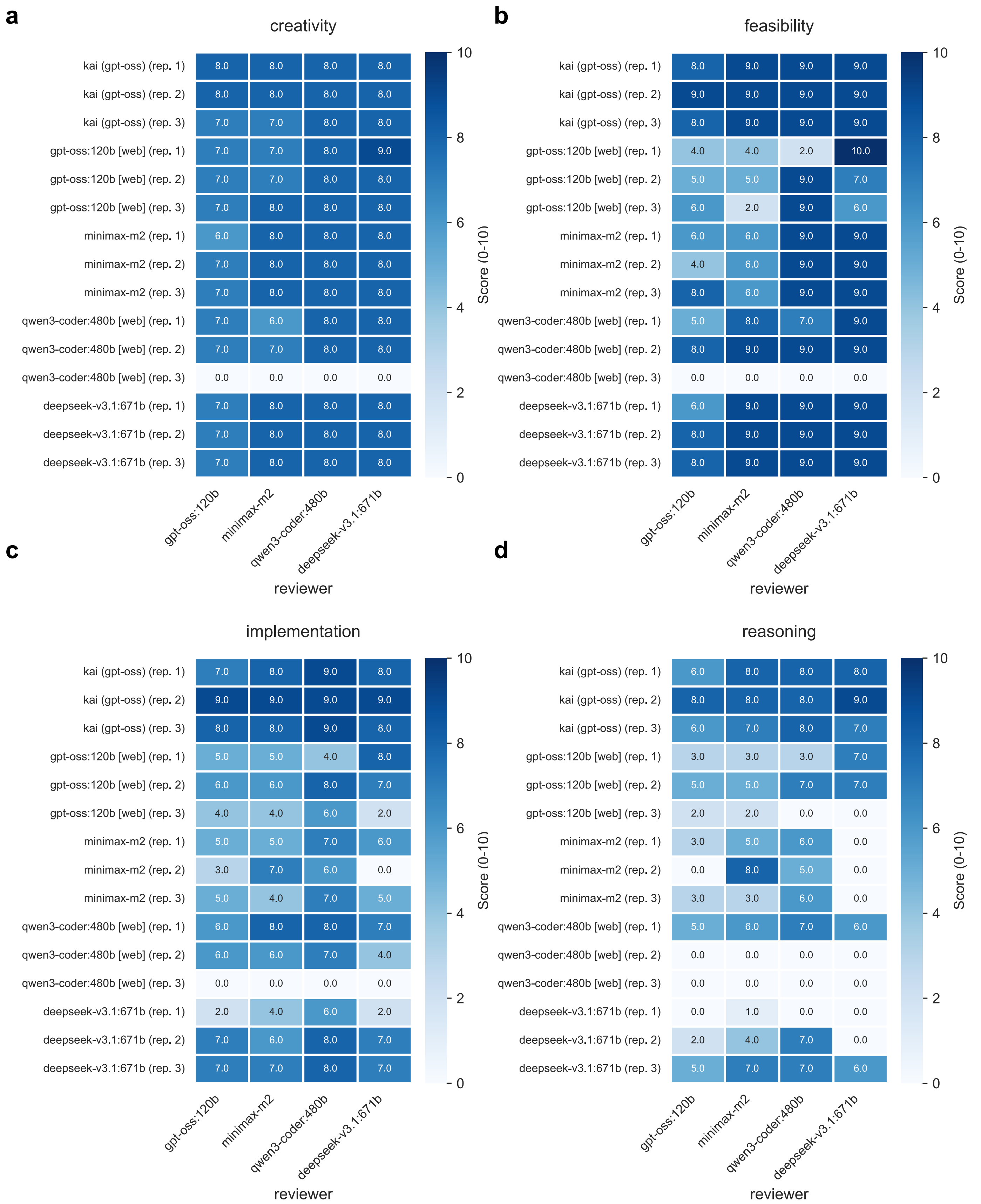


**Supp. Fig. 4: Score comparison by reviewer LLM in the open-ended scenario.** This figure extends **Fig. 3a** by showing the scores assigned by different reviewers to individual model runs. The scores creativity (**a**), feasibility (**b**), implementation (**c**), and reasoning (**d**), match the scores presented in **Fig. 3a**, the reviewers are either based *on gpt-oss:120b*, *minimax-m2*, *qwen3-coder:480b*, or *deepseek-v3.1:671b.*

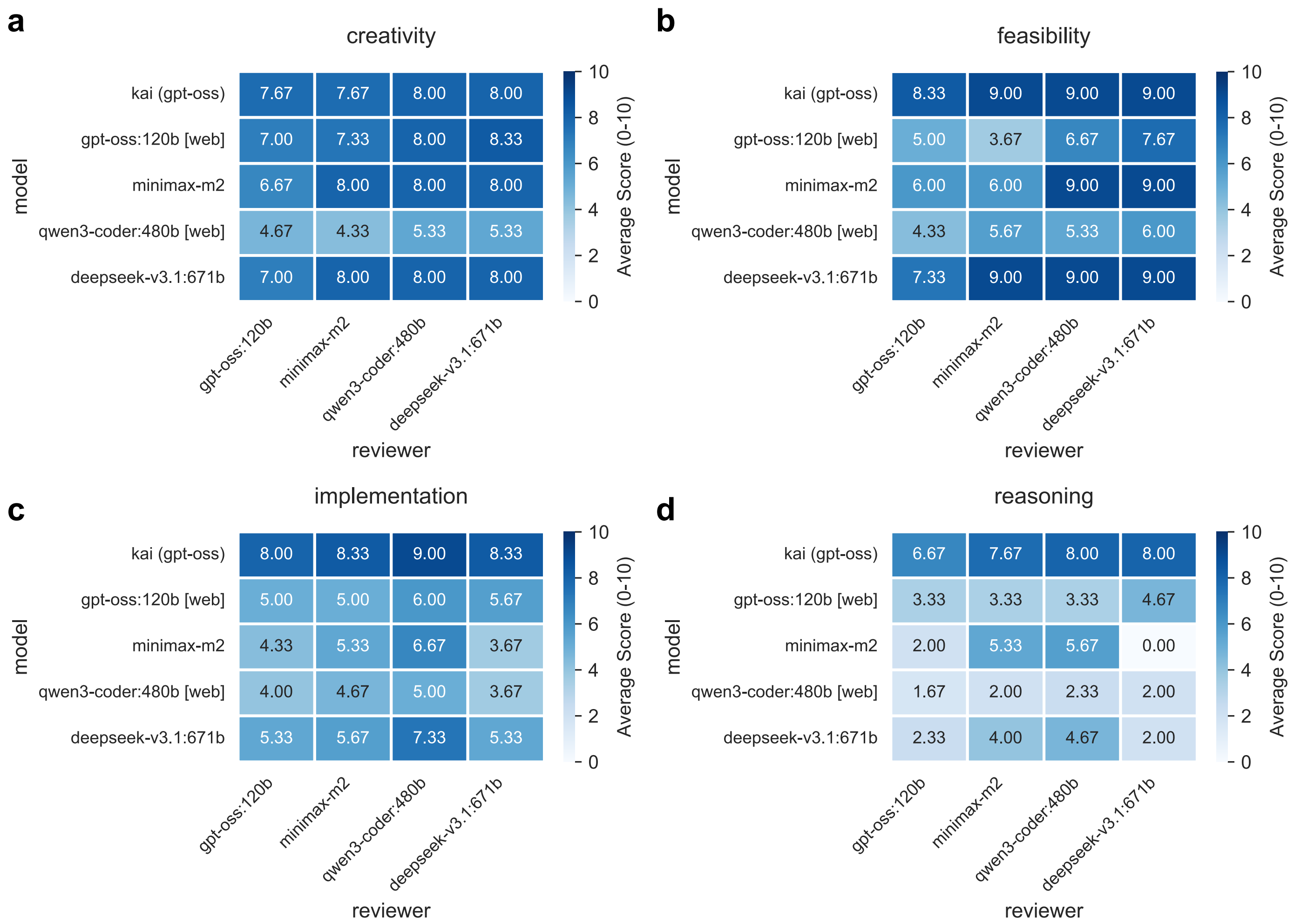


**Supp. Fig. 5: Average score comparison by reviewer LLM in the open-ended scenario.** This figure extends **Fig. 3a** by showing the average scores assigned by different reviewers to the runs of the same model. The scores creativity (**a**), feasibility (**b**), implementation (**c**), and reasoning (**d**), match the scores presented in **Fig. 3a**, the reviewers are either based *on gpt-oss:120b*, *minimax-m2*, *qwen3-coder:480b*, or *deepseek-v3.1:671b.*
